# Cross-species comparison of human and rodent primary reward consumption under budget constraints

**DOI:** 10.1101/2021.07.28.454138

**Authors:** Yue Hu, Felix Jan Nitsch, Marijn van Wingerden, Tobias Kalenscher

## Abstract

Optimal use of resources is of great importance for survival of all species. Previous studies have shown that rats optimize their allocation of a limited number of operant responses (akin to a finite budget) to trade for food rewards. Here, we propose a novel human cost-benefit decision paradigm translated from an economic task initially developed for rodents, to conduct a cross-species comparison of primary reward consumption under budget constraints. Participants had a limited budget of effortful button-presses to obtain the opportunity to drink milk rewards or watch erotic pictures. Like rodents, they adapted their choice allocation when the price (i.e. fixed-ratio requirement for reward) and the budget changed when purchasing milk rewards. However, this was not the case when purchasing the opportunity to watch erotic pictures: compared to the milk and to the rodent task, participants made more internally inconsistent, hence, irrational choices, despite otherwise identical task structures. We found non-linear changes in arousal as a function of reward selection that might underlie the irrationality in the picture task. Our study shows that the type of reward matters not only for capturing the convergence in human and non-human economic decision-making, but also for understanding the factors behind compromised economic rationality.

## Introduction

We live in a competitive world of limited resources. The principle of efficient cost allocation not only matters for species’ survival and successful Darwinian fitness maximization, it also lies at the heart of economic choice theory where an optimal decision-making strategy is crucial to maximize utility. Previous research in psychology and neuroscience focused on simple cost-benefit trade-offs, whereas in economic theory as well as in the real world, people’s demand for rewards is not only influenced by their costs, but is also constrained by the available resources at their disposal (i.e. individual income or budget). Existing literature in microeconomics has extensively discussed how limited resources affect consumers’ purchasing decisions [1-4]. Notably, animal experiments, inspired by economic demand theory, have revealed that rodents [5-8] and monkeys [9] allocate resources adaptively and optimally in response to changes in price and budget constraints. For example, rat consumers reduced their demand for a commodity if its price increases, but the reduction in demand, e.g., the demand elasticity, was less pronounced if their income (i.e. budget; here: total number of operant responses in a fixed-ratio schedule) was simultaneously increased as well - reminiscent of the effect of inflation-adjusted income on spending behavior [5, 7, 8]. Those findings showed that demand theory does not only apply to human consumers, but it is suitable to describe and predict non-human animal economic choice behavior, too. This suggests the non-trivial possibility that animals have the necessary cognitive capability to process and integrate multi-level information (e.g. prices, reward values, budget constraints) to guide an optimal use of limited resources. The fact that economic demand theory is successful in describing both human and non-human economic choice behavior demonstrates its translational face validity, but it by no means implies that humans and animals use the *same* cognitive decision processes to solve economic choice problems. What is needed for a sound cross-species comparison is a paradigm that keeps the core experimental features constant across species. Establishing such approach is not only essential for understanding complex decision making from an evolutionary perspective, but also beneficial for complementing and advancing mechanistic neurobiological models of economic choice [10-14].

To this end, we developed a novel human budget-constrained consumption paradigm that shared the most important characteristics of our rat task [5, 8]. In this task, participants made budget-constrained cost-benefit decisions to obtain access to limited amounts of rewards. To minimize procedural differences between the rat and human versions, we controlled two critical parameters: (i) price (i.e., effort requirement) and (ii) commodity (i.e., primary reinforcers). In the rodent task, animals were trained to lever-press [6] or nose-poke [5, 8] to obtain milk rewards with variable prices (i.e. the required number of operants to obtain a fixed unit of reward) and variable budgets (i.e., the total amount of operants available in an experimental session to obtain rewards). In our human version, participants had to engage in a variable number of effortful presses to obtain rewards. Importantly, while the standard microeconomic literature typically operates with monetary costs and secondary reinforcers, our rat version used milk rewards, i.e., primary reinforcers. To parallel the nature of the incentives across tasks and species, we opted for primary reinforcers in our human task as well. More specifically, inspired by previous work on economic choices with primary reinforcers [15-19], we used milk rewards and the opportunity to watch erotic pictures in separate variants of our task. The use of different incentive types allowed us to explore whether and how the kind of commodity matters for human economic decision making [20-29]. By unifying the effort and reward format as well as utilizing a similar experimental structure as in the rat task, we aimed to conduct reliable cross-species comparisons, and to evaluate the commonalities and differences in rat versus human economic decision-making.

We first present a replication of our previous findings [5, 8] on the budget-dependent effort allocation and choice adaptation in rodents (Rat Experiment). Then, we illustrate the key features of the human budget task (Human Experiment) and compare our participants’ choice behavior with that of rat consumers. This comparison revealed that participants responded to price and budget changes in a similar way to rats, but only when purchasing milk rewards, not when purchasing the opportunity to watch picture rewards. We further set out to explore the underlying mechanism and found a state-choice-rationality interaction that might drive the behavioral differences when humans traded effort budget for picture and milk rewards.

## Methods

### Rat Experiment

#### Subjects and Rewards

A total number of n = 17 male rats from two batches (batch 1: n = 7, batch 2: n = 10) were included. All rats served as control animals in two separate neurotoxic lesion experiments whose results have not been reported before (electronic supplementary material). Two flavored soymilks (i.e. chocolate and vanilla) were prepared and further diluted with water (chocolate milk, 2:3 water; vanilla milk, 1:3 water).

#### Apparatus

Animals were trained in an operant chamber (28 × 23 × 23 cm; Med Associates Inc., Georgia, VM, USA) under red light condition. The front panel of each chamber was equipped with three nose poke units arranged horizontally. All nose poke holes contained photo beams that were able to detect and signal when rats entered the hole with their snouts. On the opposite side of the chamber, the back panel was equipped with one central house light, two reward bottle access holes with infrared photo beams to detect head entry as well as two reward lights located above each access hole. Once the animal met the respective fixed ratio requirement (i.e. a given number of nose pokes), one (forced-choice trials) or two (free-choice trials) reward lights were turned on, and motorized drives lowered the reward dispenser bottle through the hole to make liquid reward accessible for two seconds when head entry into the reward port was detected. The off and on of the house light signals the beginning and ending (i.e. budget went out) of the daily session. Before the sham-lesion surgery, rats were trained in the Skinner box through five steps of shaping procedure with an increased complexity to learn the full structure of the task (see [8], for details). All inputs, outputs, and events were timestamped, controlled, and recorded by the software MedPC (Med Associates Inc., Georgia, VM, USA), and stored for offline analysis.

#### Budget and Price Conditions

The structure of the rat budget task was identical to our previous studies [5, 8] and is illustrated in Figure 1A. Rats were trained to choose between vanilla and chocolate milk rewards. They traded instrumental effort (henceforth referred to as the price of a reward, i.e. the number of nose pokes, NPs, necessary to obtain a reward) for rewards while their budget (the total number of NPs available per session that could be distributed between two rewards) was limited. In the baseline condition, each reward had a price of 2 NPs, and animals had a fixed budget of n = 80 NPs they could spend on vanilla or chocolate reward. We measured the baseline demand for chocolate and vanilla, given this price and this limited budget. Next, we changed the price of vanilla and chocolate milk to 1 and 4 NPs, respectively. Importantly, we also manipulated the budget. In the uncompensated budget condition, the budget remained at 80 NPs after implementing the price changes; that is, rats had to complete 80 NPs, as in the baseline sessions. The chocolate and vanilla prices in the compensated budget condition were identical to the prices in the uncompensated condition, but the budget was compensated following Slutsky’s demand equation [30] so that rats could, in theory, re-obtain the bundle of rewards they obtained at the preceding baseline. The budget compensation was done independently for each rat, based on their individual baseline choices. For example, assume that a rat spent its 80 NPs at baseline to purchase 25 units of chocolate and 15 units of vanilla.

**Fig.1.**
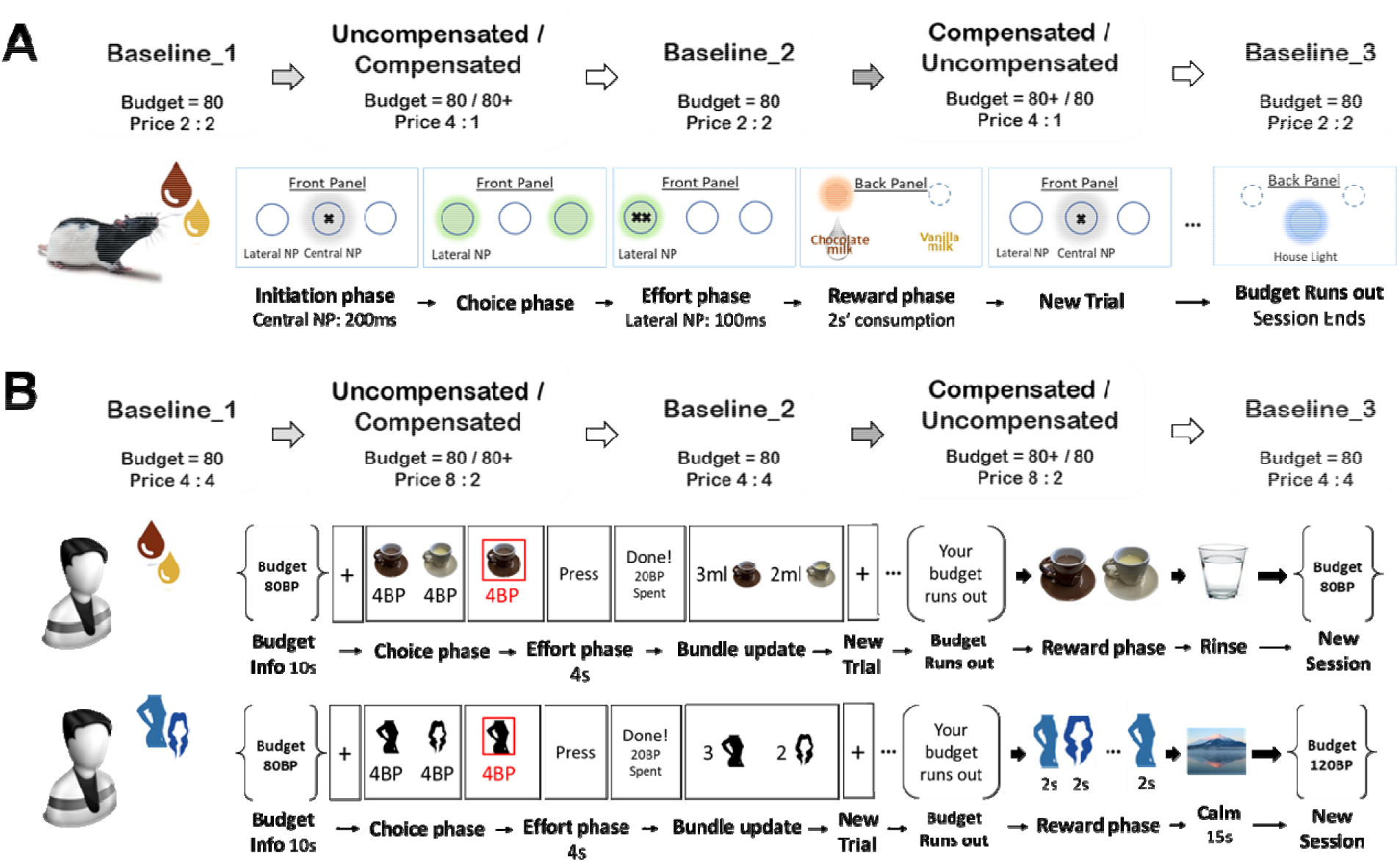
Session structure and schematic illustration of a single trial of Rat Experiment (A) and the milk and picture task of Human Experiment (B). (A) Rats initiated a trial by making a single nose poke (NP) to the activated central hole (grey color), followed by the activation of both lateral NP holes (green color). The effort requirement of each milk is indicated by the number of crosses. Daily session was terminated when the budget was used up, indicated by the illumination of the house light (blue color). The flow chart illustrates the effort requirement and budget size of each session. See Methods for details. (B) Humans participated in two similar tasks in which they made pedal presses (PP) with fingers to obtain milk rewards or the opportunity to view picture rewards. The price and budget manipulations were identical to the rat experiment. Both rewards were delivered and consumed after each session ended. See Methods for details.

To keep purchasing this choice bundle after changing the price of chocolate to 4 NPs, and the price of vanilla to 1 NP, the budget would have to be increased to 115 NPs in the compensated budget condition (25 × 4 + 15 × 1 = 115) to compensate for the price changes. That is, with a new budget of 115 NPs, the rat could purchase the same reward bundle obtained in baselines, despite the change in reward prices.

Rats went through five conditions following an A-B-A-B’-A block design, so that each price-budget condition (B or B’) was preceded and followed by a baseline condition (A). For each experimental condition, rats were trained in five continuous daily sessions to gradually learn the price and budget. We took the averaged choice data from the last two sessions of each condition, assuming that the rats’ choice patterns reached a stable state [5, 8]. The order of the two budget conditions was pseudorandomized and counterbalanced across rats. Before testing in the final task, and before the sham surgery, rats were pre-trained in a five-step shaping procedure [5]. Rats were allowed to recover for two weeks after surgery before proceeding to the final task.

#### Trial Structure

Each trial began with the activation of the central NP hole on the front panel, indicated by illumination. Rats initiated the trial by making a single NP for 200ms, followed by the activation and illumination of one (in forced-choice trials) or both (in free-choice trials) lateral NP holes. Rats were required to meet a condition- and choice-specific fixed ratio requirement (a variable number of 100ms NP responses; see Budget and Price Conditions above) to obtain a chocolate or vanilla reward. NP responses in one of the two lateral holes yielded chocolate milk reward, delivered in the back panel, responses in the other lateral hole yielded vanilla milk rewards. Rewards were presented for 2s, followed by an intertrial-interval (ITI) of variable duration such that each trial lasted exactly 32 seconds. Central and lateral NP holes were activated for a maximum period of 10s, or until rats responded. Failure to respond within 10s was followed by ITI and a start of a new trial, i.e., re-activation of the central NP hole. The NP side to reward type assignment was counterbalanced across sessions for each rat. Each daily session always started with eight forced-choice trials, four on each side in a pseudorandom order, to allow animals to sample the side-reward assignment and associated ratio requirements, followed by a variable number of free-choice trials. The forced-choice trials were not counted into the available budget in each session; i.e., rats could spend their full session-budget after having completed the forced-choice trials. A session was terminated until the animals spent the entire budget, or after 60 minutes.

### Human Experiments

#### Participants and Stimuli

Heterosexual male participants (n = 60, mean age = 24, SD = 3) were recruited. For the milk task, the same soymilks (i.e. chocolate and vanilla) that were used in the rat experiments were prepared and served in separate cups to the participants. Both flavors contained a comparable level of calories per 100ml (chocolate: 61 kcal, vanilla: 57 kcal). Prior to the milk task, participants were informed not to drink sugared beverages or eat solid food for four hours. Self-reported score (1-10, 1- not at all, 10- very much) of hunger (“How hungry are you at this moment?”) and thirst (“How thirsty are you in this moment?”) was collected before and after the milk task. Testing of milk task was only arranged around lunch time or in the late afternoon. For the picture task, two categories of pictures containing different attractive elements (attractive faces, “face”; or mildly erotic bikini/underwear motives, “body”) were prepared with a uniform size of 300 × 400 pixels. All pictures presented in the experiments were retrieved from the internet under open license. Participants needed to online rate all pictures for enjoyableness before being invited to join the laboratory part. Those who rated at least one third of pictures as highly enjoyable were selected to participate in the laboratory tasks. These selection criteria were meant to avoid participants holding an overly strong preference for one category of pictures in the experiment. Participants were also asked to try to refrain from sexual activity one day before the picture task. Self-reported score (1-10, 1- not at all, 10- very much) of sexual excitement (“How strong is your sexual desire at the moment?”) and the degree of “like to see” the pictures (“How much, in general, do you enjoy watching pictures of women with an aesthetic or sexual motive?”) was collected before and after the picture task. Each part of the experiment (online rating, picture task and milk task) was separated by at least three days to minimize any learning or carryover effects.

#### Apparatus

We opted for instrumental effortful presses [31-33] because a) effortful presses are discrete operant responses similar to the quantifiable instrumental effort requirements of the rat version of our task, and b) they allow a precise manipulations of the price of reward (i.e. number of presses) and the size of the individual budget (i.e. total amount of presses). The instrumental effort in the human experiment was implemented with a pedal press (PP) using fingers, conceptualized as equal to a button press.

#### Budget and Price Conditions

Human participants came on two separate days, with a minimum three-day interval, for the milk and picture task (Figure 1B). Participants were asked to do two tasks: the milk task and the picture task. In both tasks, participants were instructed to trade instrumental effort (here: the number of finger presses on a pedal, PPs; again referred to as the price of a reward) to obtain two types of rewards: milk rewards in the milk task (the same milk used in the rat experiments: chocolate vs. vanilla flavors) or picture rewards in the picture task (the opportunity to watch pictures of “face” and “body”). Note that in the milk task, participants obtained either chocolate-flavored or vanilla-flavored milks while in the picture task, they viewed different pictures containing one of the two attractive elements (“face” or “body”) for each selection. For each task, participants completed the five experimental conditions following a following an A-B-A-B’-A block design similar to the animal task, one session per condition, at one time. Briefly, the first, third and last condition was always the baseline condition, in which reward commodities were equally priced at 4 PPs, and individual budget was fixed at the size of 80 PPs. Between baselines were the uncompensated or compensated budget conditions in which the preferred commodity in the preceding baseline became more expensive with a price of 8 PPs and the price of non-preferred commodity was reduced to 2 PPs. We also manipulated the budget for each budget condition based on the same policy of the rat experiments. In the uncompensated budget condition, the budget remained at 80 PPs. In the compensated budget condition, the individual budget was extended following the Slutsky equation [30] to theoretically maintain participants’ baseline consumption level under the new price regime. The order of the uncompensated and compensated conditions, as well as the picture and milk task, were pseudorandomized and counterbalanced across participants.

#### Trial Structure

##### Practice sessions

Two short practice sessions with an equal price for the two commodities (first practice: 2:2 PP; second practice: 8:8 PP) were implemented before the formal experiment of each task, serving two purposes: i) to let participants get familiar with the trial structure and experience how budget was consumed, ii) to allow participants to practice on the highest level of effort in the formal task. Note that in the shaping procedure of the rat experiment we also trained rats to complete a 4:4 condition before the formal experiment.

##### Picture task

at the beginning of each condition (i.e. session), one sentence (e.g. “Your budget: 80 PPs”) was presented for 10 seconds, indicating the size of budget for this session. Each trial started with a fixation cross on the screen, followed by the appearance of two icons representing two categories of the picture rewards with each price (e.g. “4 PP”) written below (Figure 1B). The side (i.e., L/R) of each category was randomized across trials. When participants made their choice, the chosen commodity was shortly highlighted and then the effort phase began. In the effort phase, participants pressed buttons to reach the required effort level within four seconds. The length of the effort phase was calibrated in a pilot study to avoid long waiting. If participants were too slow in making their choices or failed to reach the effort requirement within the given time, this trial was considered as missed trial and restarted. When the effort requirement was met, and four seconds had passed, participants received an update of their budget expenditure and the accumulated choice selection of rewards (i.e. updated choice bundle). When the budget expired, participants received the notification at the end of the last trial and proceeded to the reward consumption phase in which picture rewards were displayed for 2 seconds in the same order as participants chose them. For example, if participant A chose to view a body picture in the first trial and then a face picture in the second and third trials, the presentation of pictures followed the same order of “body picture 1 - face picture 1 - face picture 2 - …”. After the consumption phase, a photo of Mount Fuji was presented for 15 seconds as a between-session cue, and then proceeded to the next experimental condition.

##### Milk task

The milk task applied the same trial structure as the picture task except that the milks were served in two cups after the session budget expired. Participants had 40 - 60 second to consume the milk rewards, which was similar to the duration of the reward consumption phase in the picture task. After consuming the rewards, they were provided with a cup of water to rinse their mouth and then proceeded to the next condition.

#### Control Experiments

To provide additional validations of the results obtained from our novel translated human paradigm, we also present two control experiments to investigate the effect of milk reward satiation and timing of picture reward consumption on participants’ behaviors (electronic supplementary material).

### Analysis

#### Demand elasticity as a measure of choice adaptation

To capture the degree by which individual choices of rewards changed in response to price and budget changes, we calculated demand elasticity as an index of price sensitivity, using linear regression on the log-transformed number-of-choices/price-ratio pairs as given by the equation ([6], see also [5, 8]):

(1)

Where *q*_*i*_ indicates the quantity of reward (i) chosen, *p*_*i*_ */p*_*j*_ indicates the price ratio for reward (i) over reward (j) per price/budget regime and α is a constant. *ε* denotes the estimated elasticity in demand. We estimated *ε* individually for preferred and non-preferred reward for the two budget conditions, separately. The negativity/positivity of *ε* represents the direction of change in demand in response to the price and/or budget changes. The absolute value of *ε* measures the extent of change in demand compared to the baseline levels. For example, a more negative *ε* parameter indicates a stronger reduction in demand for rewards after a price increase, or a larger increase in the demand for rewards after a price decrease. A positive *ε* indicates an increased demand for a more expensive reward, or a reduced demand for a cheaper reward. Economic demand theory predicts a general negative *ε* if prices go up, indicative of a reduction in demand, and vice versa if prices drop.

#### Choice consistency as a measure of decision rationality

Our research design offers the possibility to quantify our participants’ degree of rationality by analyzing their compliance to consistency criteria in revealed preference theory (Generalized Axiom of Revealed Preference; GARP [34-36]). In brief, GARP requires that participants make internally consistent choices. For example, choice chains should be transitive (if option A is chosen over option B, and B is chosen over C, A should also be chosen over; for details on GARP and consistency, refer to [37, 38]). To infer individuals’ degree of decision rationality from their choice data, accumulated over each experimental condition, we counted how many choice bundles posed a violation of GARP. For participants who failed to maximize their utility of consumption, we should observe some degree of GARP violations.

#### Statistical analysis

Choice and elasticity data were analyzed in JASP [39] using repeated measures ANOVA with Bonferroni correction for multiple post-hoc comparisons. Wilcoxon signed rank test was performed in R for comparing the count of GARP violation between the picture and milk task, as well as for comparing self-reported score of state before and after each task. The linear modeling on the counted data of GARP violation against the continuous variable (cumulative choice of preferred rewards) and the categorical variable (picture vs. milk task) was implanted in a general Poisson linear regression (*glm* function from MASS package, [40]). The regression on the score of post-experiment sexual excitement against the summed reward intake was also estimated by a hyperbolic function with the *glm* function. The raincloud plots on self-reported state were created with ‘raincloud plot’ R package [41].

## Results

### Cross-species comparison of the budget effect on reward elasticities

#### Participants showed a rigid preference when purchasing preferred picture rewards

First, we checked whether manipulating budget and price influenced budget and choice allocations in rats and human participants (Figure 2A-C). A 2 (batch) x 5 (condition) mixed ANOVA on the rats’ choices of preferred rewards (which was, typically, chocolate) revealed no significant interaction between the animal batch and condition (F_4, 75_ = 0.594, p = 0.614, partial η^2^ = 0.038), therefore, we combined the two batches together for further analysis.

**Figure 2.**
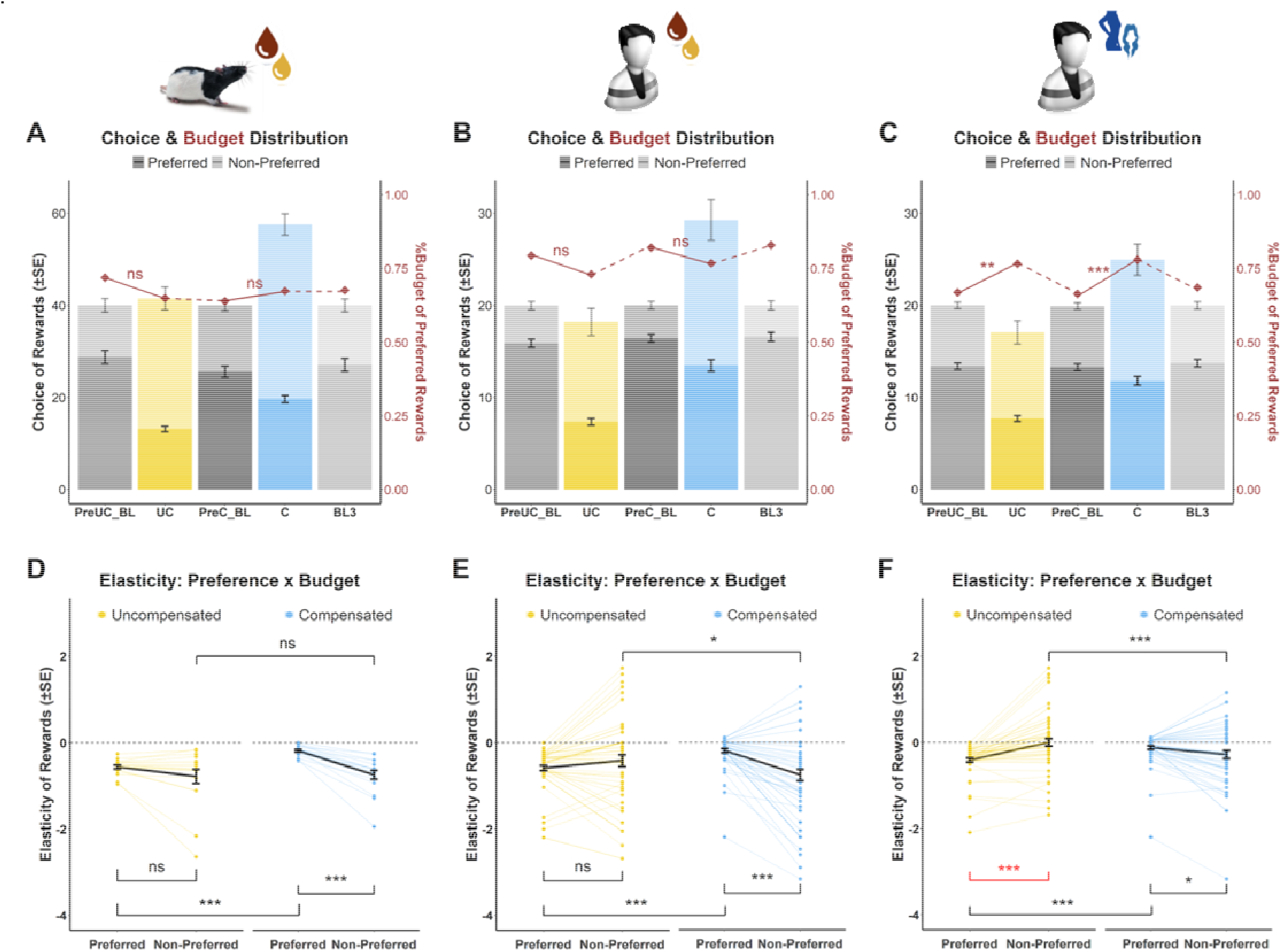
Participants behaved in a similar manner to rat consumers in the milk task, but not in the picture task. (A-C) Stacked bar plots of choice distribution on preferred and non-preferred rewards (left y-axis) and averaged proportion of budget spent on preferred rewards (right y-axis) across five experimental conditions for rat consumers (A) and participants in the milk (B) and picture (C) task. Participants in the picture task significantly increased their budget spent on expensive commodities in both budget conditions, showing a rigid preference for preferred picture rewards. UC: uncompensated budget condition; C: compensated budget condition; PreUC_BL: preceding baseline of uncompensated budget condition; PreC_BL: preceding baseline of compensated budget condition; BL3: the third baseline condition. (D-E) Demand elasticity of preferred and non-preferred rewards in the uncompensated and compensated budget of rat consumers (D) and participants in the milk (E) and picture (F) task. ***: p < 0.001; **: p < 0.01; *: p < 0.05; ns: not significant.

Consistent with our previous experiments [5, 8], rats reduced their demand for rewards that became more expensive, and increased their demand for rewards that became cheaper (all p < 0.001 for comparisons of chocolate purchases in uncompensated vs. compensated and [uncompensated/compensated] vs. baselines, paired t-tests) while their reward preferences remained stable across three baseline conditions (all p > 0.103, paired t-test, Figure 2A). For the human experiment (Figure 2B-C), we ran a 2 (task: picture vs. milk) x 5 (condition) ANOVA on the choices of preferred reward. We found a main effect of task (F_1, 59_ = 21.111, p < 0.001, partial η^2^ = 0.077) and condition (F_4, 236_ = 159.306, p < 0.001, partial η^2^ = 0.415) on choices, as well as a significant interaction of *task x condition* (F_4, 236_ = 8.618, p < 0.001, partial η^2^ = 0.033). In both tasks, human participants significantly reduced their demand of preferred reward when prices increased (all p < 0.002 for uncompensated vs. compensated and [uncompensated/compensated] vs. baselines, paired t-tests). They also exhibited a higher baseline selection of preferred reward in the milk task than the picture task (all p < 0.001 for the three baselines, paired t-test). Together, these revealed a consistent price effect on demand across species and reward tasks.

Interestingly, given the reduced demand for preferred commodities in the picture task, the proportion of budget that participants spent on purchasing preferred rewards was much higher than the preceding baseline in both budget conditions (uncompensated vs. baseline: t = 2.977, p = 0.004, Cohen’s d = 0.384, 95% CI [0.121, 0.645]; compensated vs. baseline: t = 3.707, p < 0.001, Cohen’s d = 0.479, 95% CI [0.209, 0.744]; paired t-tests, Figure 2C). In other words, participants’ genuine preference for preferred rewards remained intact after the price increase in the picture task. Such a significant shift in the budget allocation was not found in the rat experiment (uncompensated vs. baseline: t = -1.671, p = 0.114, Cohen’s d = -0.405, 95% CI [-0.895, 0.096]; compensated vs. baseline: t = 1.054, p = 0.307, Cohen’s d = 0.256, 95% CI [-0.232, 0.735]; pairwise t-tests) or the human version of milk task (uncompensated vs. baseline: t = - 1.585, p = 0.118, Cohen’s d = -0.205, 95% CI [-0.459, 0.052]; compensated vs. baseline: t = -1.51, p = 0.136, Cohen’s d = -0.195, 95% CI [-0.45, 0.061]; pairwise t-tests, Figure 2A-B). These results indicate a rigid preference for picture rewards when spending limited budget to purchase priced commodities.

#### Rat and human consumers exhibit similar reward elasticity patterns in the milk task

We, next, determined if the choices of our rat and human consumers only depended on reward prices, or, additionally, also on the budget size, as would be expected by budget effects on demand mentioned above [5, 8]. Budget effects on choice are puzzling since they imply that subjective costs of effort (here: NPs in rats or PPs in humans) is not equivalent to the net effort, but depends on the available budget: 4 NPs are less aversive if a rat has a large budget to spend. To see if budget mattered, we quantified the degree by which demand elasticity, as a proxy for change in demand, depended on reward preference, reward price and budget. For the Rat Experiment, we conducted a 2 (budget: compensated vs uncompensated) x 2 (preference: preferred vs non-preferred) ANOVA on demand elasticity (Figure 2D). We found a main effect of preference (F_1,16_ = 25.381, p < 0.001, partial η^2^ = 0.613) and a significant *budget x preference* interaction (F_1,16_ = 5.619, p = 0.031, partial η^2^ = 0.26). In the uncompensated budget condition where the budget remained the same as in the preceding baseline, most of the rat consumers reduced their selection of preferred, but now more expensive milk rewards (i.e. price increase from 2 NPs to 4 NPs), and increased their selection of the non-preferred, now cheaper non-preferred rewards (priced dropped from 2 NPs to 1NP); the comparison of elasticities of the two rewards revealed no significant difference (t = 1.744, p = 0.1, Cohen’s d = 0.423, 95% CI [-0.08, 0.914]), suggesting that demand for the preferred reward decreased in response to the price increase as much as demand for the non-preferred reward increased in response to the price drop. In the compensated budget condition where rats had a larger budget of NPs to spend, they selected more of the preferred, expensive reward compared to the uncompensated budget condition, thus, exhibiting a less negative demand elasticity of preferred reward in the compensated compared to the uncompensated condition (t = -8.935, p < 0.001, Cohen’s d = -2.167, 95% CI [-3.04, 1.274]). Such behavioral shifts in response to the budget extension also enlarged the difference between the two reward elasticities in the compensated budget condition (t = 6.86, p < 0.001, Cohen’s d = 1.664, 95% CI [0.91, 2.396]), but not in the uncompensated condition (t = -0.205, p = 0.84, Cohen’s d = -0.05, 95% CI [-0.08, 0.914]). Thus, we replicated our previous findings on the budget-dependent choice adaptation in rats [5, 8].

To test whether human participants exhibited similar budget effect on demand elasticity, we performed a three-way ANOVA with budget condition (uncompensated, compensated), task (milk, picture) and preference (preferred commodity, non-preferred commodity) on reward elasticity. We found a main effect of reward type (F_1,59_ = 11.240, p = 0.001, partial η^2^ = 0.16), suggesting that participants’ demand for milk rewards was much more elastic to the price changes than their demand for picture rewards. We also found a significant *budget x preference* interaction (F_1,59_ = 62.551, p < 0.001, partial η^2^ = 0.515) and a significant interaction of *preference x task* on demand elasticity (F_1,59_ = 14.668, p < 0.001, partial η^2^ = 0.199, electronic supplementary material, Figure S1). The three-way interaction of *budget x preference x task* was not significant (F_1,59_ = 1.557, p = 0.217, partial η^2^ = 0.026). Therefore, we performed separate 2 (budget: compensated vs. uncompensated) x 2 (preference: preferred vs. non-preferred) ANOVAs on reward elasticity for the milk and picture tasks. The picture results will be described further below.

In the milk task (Figure 2E), we found a main effect of preference (F_1,59_ = 5.892, p = 0.018, eta square = 0.091) and a significant interaction of *budget x preference* (F_1,59_ = 31.786, p < 0.001, partial η^2^ = 0.35) on reward elasticity. This suggests that, similar to the rat consumers, participants adapted their choice of milk rewards when purchasing from a constrained budget (i.e. uncompensated budget condition) and their elasticity for the preferred milk reward did not differ from the non-preferred milk reward (t = -1.668, p = 0.101, Cohen’s d = -0.215, 95% CI [-0.47, 0.042]). Like with rats, the comparison was significant in the compensated budget condition, showing that the elasticity of non-preferred milk was much more negative than the elasticity of preferred milk (t = -5.325, p < 0.001, Cohen’s d = -0.687, 95% CI [-0.967, -0.403]). Our participants’ elasticity of preferred milk in the compensated budget condition was also much less negative than that of the uncompensated budget condition (t = 7.152, p < 0.001, Cohen’s d = 0.923, 95% CI [0.618, 1.223]). In addition, their demand for non-preferred reward in the compensated budget condition was more elastic than in the uncompensated condition, too (t = -2.316, p = 0.024, Cohen’s d = -0.299, 95% CI [-0.556, -0.039]). Taken together, the choice and elasticity data in humans replicate our previous findings on the price and budget effect on rats’ consumption behaviors and further reveal a cross-species similarity in the elasticity pattern when trading effort for milk rewards with budget constraints.

#### Picture task: participants’ rigid preferences leads to a non-negative elasticity of demand

The elasticity results in the human milk task suggest a budget-dependent choice behavior when trading effort for milk rewards, just like we observed in the rat task. Both humans and rats tended to reduce their demands for expensive and preferred milk rewards when the budget cap was tight, and relaxed their restricted selection of priced milk rewards when a larger budget was available. In the uncompensated budget condition where the individual budget remained fixed (i.e. 80), the total amount of commodities that individuals can afford (i.e. purchasing power) decreases because of the changes in reward prices. Hence, a price-sensitive consumer should select more of the cheaper rewards to maintain an adequate amount of reward intake, which would result in an increased negative elasticity of demand for the two reward commodities, a phenomenon observed in our rat consumers [5, 8], and in the human milk task. However, this expected shift to the cheaper option was not found in the picture task. To examine the effect of price and budget on reward elasticity in the picture task more closely, we ran the same analysis as above in the picture task and found a main effect of preference (F_1,59_ = 5.853, p = 0.019, partial η^2^ = 0.09) as well as a significant interaction of *budget x preference* (F_1,59_ = 61.552, p < 0.001, partial η^2^ = 0.511) on the elasticity of demand for picture rewards (Figure 2F). In line with our observation and in contrast to the rats and humans in the milk task, the elasticity of non-preferred picture reward was much more positive than the elasticity of preferred reward in the uncompensated budget condition (t = 6.737, p < 0.001, Cohen’s d = 0.87, 95% CI [0.569, 1.165]). This suggests that, unlike in the milk task, participants in the picture task *reduced* their selection of non-preferred picture rewards after its price decreased, thus showing a reversed price-effect on demand. Interestingly, we found that this pattern was budget-dependent, as the elasticity of non-preferred reward was much less positive in the compensated budget condition than the uncompensated condition (t = -3.295, p = 0.002, Cohen’s d = - 0.425, 95% CI [-0.688, -0.159]). These findings indicate that the reversed price effect, i.e., the decrease in demand with decreasing prices, was attenuated when having an extended budget to spend. In addition, the elasticity of non-preferred picture rewards was more negative than the elasticity of preferred rewards in the compensated budget condition (t = -2.401, p = 0.019, Cohen’s d = -0.31, 95% CI [-0.568, -0.05]). The elasticities of preferred picture reward significantly differed between the two budget conditions, too (t = -7.33, p < 0.001, Cohen’s d = - 0.946, 95% CI [-1.249, -0.638]). Thus, crucially, the cross-species similarity in choice behavior when trading efforts for rewards was confined to milk rewards; when trading efforts for picture rewards, participants exhibited a reversed price effect on demand: their demand for the non-preferred pictures *decreased* when their price dropped, and their budget spent on preferred pictures largely increased to maintain an unchanged demand when their price increased.

### Within-subject comparisons in the milk and picture tasks

#### Higher choice consistency in the milk task than the picture task

We, next, counted how many choice bundles, out of the five bundles generated in each session, violated GARP to quantify if the reversed price effect on demand in the picture task compromised rationality, compared to the milk task. To satisfy GARP, individuals must reveal stable preferences in their decisions, as well as act cost efficiently in the pursuit of these [37, 38]. In other words, GARP is satisfied when a participant does not show circularity (e.g. if their choices meet transitivity requirements, see above) in the selected choice bundles. If drawing the chosen bundles along the budget lines, one bundle is at least as good as other bundles on the same budget line, and better than bundles that locate underneath its budget line. For example, among the five choice bundles of participant 27 in the picture task (Figure 3B), bundle “BL3” is as good as bundle “preUC_BL” and “preC_BL”, and bundle “C” is better than all other four bundles. Bundle “UC”, underneath the budget line of baseline (the shadow triangle), is worse than the three baseline bundles. However, bundle “BL3” is also not better than bundle “UC” since it locates below the “UC” budget line. Thus, the bundle “UC” and “BL3” constitute two violations of GARP. We found a significant difference in the violation count between the milk and the picture tasks (p = 0.013, Wilcoxon signed-rank test, Figure 3A). This indicates that human participants behaved less rationally when consuming picture rewards than milk rewards.

**Figure 3.**
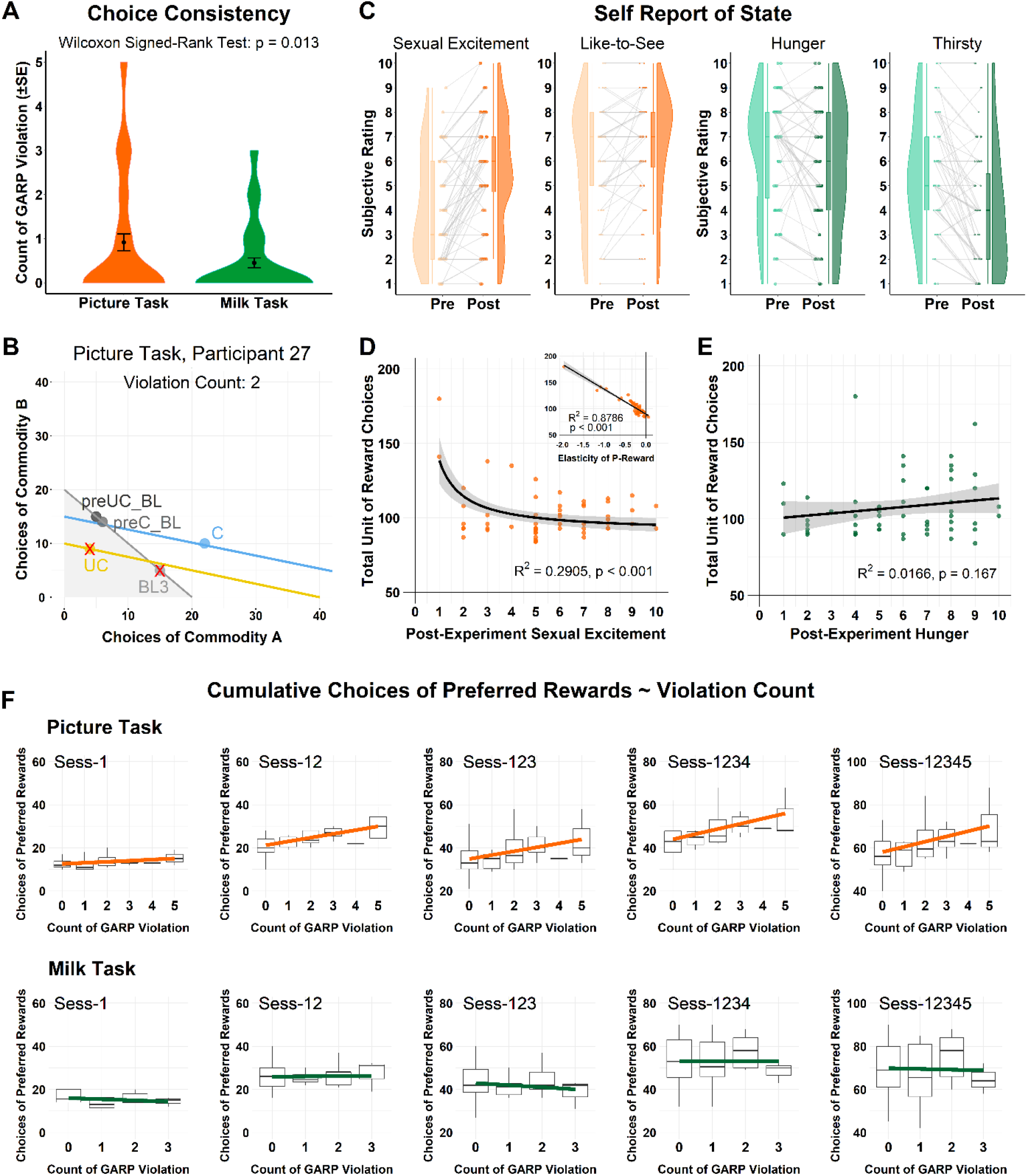
Choice-state-rationality in the picture and milk task of Human Experiment. (A) Count of GARP violation, as an index of choice consistency, in the picture and milk task. (B) An example of individual choices generated in the five experimental sessions of the picture task. Red cross indicates the two choice bundles that violate GARP. (C) Self-reported scores before (Pre) and after (Post) the task. (D) Hyperbolic curve fitting on post sexual excitement and total amount of reward intake in the picture task. The small figure on the top right plots the linear correlation between elasticity of preferred rewards (P-reward) and total amount of reward intake. (E) Linear regression on post hunger against the total amount of reward intake in the milk task. (F) Boxplots of the cumulative consumption of preferred rewards against the level of GARP violation across five conditions (i.e. session) in the picture and milk task. preUC_BL: baseline preceding the uncompensated budget condition; preC_BL: baseline preceding the compensated budget condition; BL3: the last baseline condition; UC: uncompensated budget condition; C: compensated budget condition.

#### Non-linear choice-state correlation in the picture task

Why did our participants show a reversed price effect on demand when trading effort for picture rewards, but not with milk rewards, despite similar experimental structures? Why were they more irrational when dealing with pictures than with milk? One probable reason for the difference in choice behavior between tasks is the nature of the primary rewards that are likely linked to different visceral states (i.e. hunger and sexual appetite). One theory [42] posits that utility maximization can be formalized as a function of visceral state (e.g., hunger) and consumption activity (e.g. milk drinking) that would modulate individuals’ marginal utility of goods and drive a consequent change in the rate of reward substitution. It also predicts that an instant change in consumption would further mitigate or elevate individual states (e.g., reduce hunger), thus, altering the marginal utility of future rewards. Applied to our task, there might exist a distinct choice-state interaction that accounts for the observed difference in the choice allocation and adaptation in the milk and picture task. For example, continued consumption of erotic pictures might change the arousal state as the task progresses, which, in turn, might affect, e.g., increase, the demand for preferred erotic pictures. We hypothesized that this cross-talk between visceral state and demand for reward was different in the picture than the milk task. Thus, we predicted that our participants’ visceral states changed after the consumption of picture and milk rewards in a reward-dependent way. We furthermore explored if, and to what degree, the amount of GARP violations changed as the milk and picture tasks progressed.

To determine the predicted co-dependency of visceral states and choices in the milk and the picture tasks, we then ran a Wilcoxon signed-rank test on paired scores (Pre/Post, Figure 3C) of subjective ratings on their state of general sexual excitement and “like to see” the pictures in the picture task, and thirst and hunger in the milk task. Participants reported a reduced hunger (t = -2.565, p = 0.013, Cohen’s d = -0.337, 95% CI [-0.6, 0.071], pairwise t-tests) and thirst (t = - 6.118, p < 0.001, Cohen’s d = -0.797, 95% CI [-1.087, 0.501], pairwise t-tests) after participating in the milk task, but an increase in sexual excitement (t = 6.765, p < 0.001, Cohen’s d = 0.873, 95% CI [0.573, 1.169], pairwise t-tests) and increased degree of “like to see” (t = 3.153, p = 0.003, Cohen’s d = 0.407, 95% CI [0.142, 0.669], pairwise t-tests) after the picture task. We found no significant choice/elasticity – state correlations in the milk task (all p > 0.167, Figure 3E). However, we observed a strong negative correlation between the post-experiment score of sexual excitement and the total unit of reward intake (i.e. bundle size, disregarding reward commodity), i.e., participants with higher arousal viewed less pictures in the task, meaning they chose more often the expensive preferred pictures. Notably, this relationship was better approximated by a hyperbolic (R-squared = 0.291, p < 0.001, AIC = 489.19, Figure 3D) than a linear function (R-squared = 0.049, p = 0.05, AIC = 505.76), indicating a non-linear or spillover development of sexual arousal with the consumption of picture rewards. In addition, the overall bundle size was linearly correlated with the mean elasticity of preferred rewards (R-squared = 0.879, p < 0.001, Figure 3D), suggesting that a more negative elasticity yielded a higher amount of reward intake.

Put together, these findings confirm our prediction on the state change in the two tasks but reveal a counterintuitive choice-state correlation in the picture task, which can be best depicted by a hyperbolic decay. Specifically, participants who exhibited an elastic demand for expensive pictures and, thus, obtained a larger bundle size of rewards, reported a less aroused state after task completion. Those participants who perseverated on selecting the preferred, but costly pictures obtained a smaller number of picture rewards in total and reported higher post-experiment arousal levels. Critically, such correlation was prominent in the participants with a low-to-medium level of arousal state and flattened among highly aroused individuals.

#### Early emerged rationality-choice interaction in the picture task, not in the milk task

Our previous analysis demonstrated that an increased arousal state correlated negatively and nonlinearly with the total consumption of picture rewards. To examine whether there exists a link between the consumption of preferred rewards with decision rationality, we ran a Poisson regression on the count of GARP violation against the summed choices of preferred picture or milk rewards. We found a significant positive correlation in the picture task (z = 3.854, p < 0.001), but not in the milk task (z = -0.226, p = 0.821). This indicates that a higher violation of GARP choice consistency was associated with a higher selection of preferred picture rewards. So far, we found a choice-state and choice-rationality correlation in the picture task, but no indication of significant correlations in the milk task. To explore where there exists a session-wise choice-rationality correlation, we performed a generalized linear model utilizing a Poisson regression on the count of GARP violations against the cumulative choices of preferred rewards, the two tasks (milk vs. picture) and the interaction between the two (Figure 3F). We found a significant interaction between the task and the cumulative choice of preferred rewards on GARP violations that emerged from the beginning of the tasks (Sess-1: z = -2.898, p = 0.004) and persisted throughout the five progressive sessions (Sess-12: z = -2.343, p = 0.019; Sess-123: z = - 2.819, p = 0.005; Sess-1234: z = -2.533, p = 0.011; Sess-12345: z = -2.254, p = 0.019). Importantly, individuals’ selection of preferred rewards in the earliest session was predictive of their count of GARP violations, and such association remained significant throughout the whole picture task (Sess-1: z = 3.08, p = 0.002; Sess-12: z = 4.836, p < 0.001; Sess-123: z = 3.895, p < 0.001; Sess-1234: z = 4.427, p < 0.001; Sess-12345, z = 3.854, p < 0.001). Notably, such prediction was not present in any sessions of the milk task (Sess-1: z = -1.372, p = 0.17, Sess-12: z = 0.192, p = 0.848, Sess-123: z = -0.941, p = 0.347, Sess-1234: z = -0.003, p = 0.998; Sess-12345, z = -0.226, p = 0.821).

Note that the individual counts of GARP violations were estimated based on their overall five choice bundles generated through the five experimental sessions of each task. The early-emerging predictive power of preferred reward selections on the choice consistency further supports and adds new evidence to the mechanistic account of irrational behaviors in the picture task. Participants generally expressed a low level of arousal at the beginning of the task (Figure 3B). Their state of arousal was gradually elevated after viewing the collected picture rewards. The elevated state might have induced a *projection bias*, a decision bias driven by a focus on the current state, thus, aroused participants failed to adapt their choices to the changing environment [42-45]. Such choice-rationality interaction was absent in the milk task across all sessions, suggesting that participants experienced a different or non-significant mechanistic regulation of state and choice on decision rationality when trading effort for the milk rewards.

To conclude, our analysis of our human data provides a mechanistic account of the irrational behaviors in the picture task. We found three pieces of evidence in supporting of a choice-state interaction that drives the irrational behavior during the picture reward consumptions: 1) individuals’ post-experimental state of arousal was negatively hyperbolically correlated to their summed collection of picture rewards; 2) a higher selection of preferred rewards was linked to a higher GARP violation; 3) decision rationality can be predicted by the choice of preferred rewards from the earliest session and throughout the whole task. Put in simple words, we believe that a positive feedback loop of arousal and picture preference can explain our human participants’ irrationality in the picture task: participants were aroused by watching their preferred pictures, which, in turn, increased their desire for watching more of their preferred pictures, regardless of its increased price, which further increased their arousal levels, along with a consequent increase in the demand for their preferred pictures.

## Discussion

Rodents reveal choice patterns and biases during economic decision making, similar to humans, including regret [46], sunk-cost sensitivity [47], risk preference [48] and inequity aversion [49]. Our previous studies also showed that rats complied well with the prediction of demand theory in making budget-dependent cost-benefit decisions [5, 8]. The central aim of the present study was to develop a parallel human paradigm translated from the rat budget task to investigate humans’ budget spending behavior and to conduct reliable cross-species comparisons.

Previous studies using primary rewards showed a common effort discounting phenomenon in rats and humans [16, 50, 51]. Here we also observed a price-sensitive choice pattern in both species, reflected by the negative elasticity of preferred rewards, i.e., a drop in demand when the price of a reward increased, and a rise in demand when the price of a reward decreased. Also, our data revealed a consistent budget-dependent effort-reward trade-off across species and reward type in which the elasticity of preferred rewards was less pronounced in the compensated than the uncompensated budget condition, replicating the budget effect on reward elasticity previously shown in rat consumers [5, 6, 8].

In line with our previous finding on the baseline-preference-dependent choice shift after the price/budget change [8], we observed a similar adaptive behavior in our rat consumers, as well as in humans in the milk task (Figure 2, Figure S2). This hints at similarities in the decision strategies employed by both species when utilizing limited budgets to obtain food rewards. Importantly, our participants’ choice pattern in the picture task was different than their choices in the milk task, or the rats’ choice in the rodent version of the milk task: participants showed a rigid preference for priced commodity and a reversed price effect on demand: participants *reduced* their selection of the non-preferred rewards when their prices dropped, and their demand for the preferred reward remained insensitive to the rise in price. This decision strategy went along with a higher violation of GARP, suggesting that they made more inconsistent, hence, irrational, choices in the picture task compared to the milk task (Figure 3, Figure S4). In addition, we included two control experiments to validate the main results of human budget experiment (Figure S1). Thus, we concluded our findings of the distinct strategy of budget and choice allocation regarding the two primary rewards to be a robust and replicable behavioral phenomenon on human participants.

Thus, by developing and running a budget task that was carefully translated from its rodent equivalent, we found a cross-species similarity in choice allocation and strategy adaptation when humans and rats spent limited budget to obtain and consume primary rewards. Crucially, this comparability only holds for the milk tasks, and not for the picture task where participants showed more inconsistent behavior. We further showed that this cross-species comparison was independent from the difference in the sample size of human and rodent experiment (Figure S3). Thus, the nature of the reward seems to matter for the degree of rationality. There have been concerns regarding a *description-experience gap* when running comparative studies on humans and non-human animals [13, 52-56]. It refers to the discrepancies between choices made in description-versus experience-based paradigms. This methodological gap raises challenges for interpreting shared decision process and/or underlying neural mechanism of similar behavioral phenomenon, for example, risk preference, identified from human and animal experiments using description- or experience-based gambling tasks. Similarly, studies on humans’ budget-related purchasing decision making were mostly limited to description-based protocols [2, 4, 57-60], with a few exceptions in which participants experienced a real-time budget-spending scenario to obtain monetary rewards [61-63]. Our study provides empirical evidence on how the consumption behaviors of humans and rats converge in a decision task involving experienced outcomes, and how the disparities in their choice pattern arise by the types of rewards being consumed.

Our analysis also provides a plausible mechanistic account of the irrationality in the picture task (Figure 3). Severe violations of GARP have previously been observed in individuals with impaired or less developed cognitive functions [64-66] or with a mood disorder [67]. These findings support the notion that individuals’ cognitive limitation or emotional state might contribute to economic irrationality [38, 42, 68, 69]. Following this line of reasoning, participants indeed reported an increased state of sexual arousal after the picture task and a reduced hunger after the milk consumptions. Importantly, the individual state was found negatively correlated to the total reward selections only in the picture task. The decision to view more of the preferred pictures, implying less picture consumptions in total, led to a higher post-experimental state of arousal. In addition, the preference strength for preferred picture rewards predicted decision rationality. Also, the predictive power of cumulative reward selection emerged already in the early sessions and persisted throughout the whole task. These findings are in line with a model [42, 43] on a mutually regulatory role of elevated state and consumption activity that affects current and future utility maximization of economic behaviors. Not having a direct observation on the development of arousal state, our behavioral analysis cannot conclusively elucidate the causality between state and choice during the reward consumptions. Future studies that measure automatic responses of arousal during the decision process (e.g. pupil dilation [70]) will provide more objective and concrete evidence to address the underlying psychological or neural mechanisms.

## Conclusion

We aimed to conduct a sound between-species comparison of economic choice behavior. Our results show that the type of consumed rewards carrying distinct innate values could play a critical role for capturing behavioral parallels in human and non-human animals. We further examined the choice-state correlation that might account for the irrationality in participants’ economic choice behaviors. Our translational budget task will allow us to investigate the psychological and neural mechanism of budget-spending behaviors in humans and rats.

## Supporting information

Electronic Supplementary Material

## Data and code sharing

The raw data and analysis scripts are publicly available on the Open Science Framework (https://osf.io/9yqbk/).

## Acknowledgements

We thank Julia Neuhaus, Marc Deibert, Melanie Mialki, Agathe Guist and Moujan Rezvani for their help with data collection. We thank Manuela Sellitto for her comments on the current version of the manuscript.

## Author contributions

Y.H.: Conceptualization, Methodology, Validation, Formal analysis, Investigation, Data curation, Software, Writing – original draft preparation, Writing – review & editing, Visualization, Project administration, Funding acquisition.

F.J.N.: Formal analysis, Writing – review & editing.

M.vW.: Writing – review & editing.

T.K.: Conceptualization, Resources, Writing – review & editing, Supervision.

## Ethics

Ethics approvals for the human experiments have been obtained from the institutional review board of the Institute of Experimental Psychology, Heinrich-Heine University Düsseldorf. All participants gave their informed consent. The study was conducted in alignment with the declaration of Helsinki. Ethics approvals of animal experiments were granted by the Landesamt für Natur und Verbraucherschutz (LANUV) of Northrhine Westfalia.

## References

1. Du R.Y., Kamakura W.A. 2008 Where did all that money go? Understanding how consumers allocate their consumption budget. Journal of Marketing 72(6), 109–131. (doi:10.1509/jmkg.72.6.109).

2. Homburg C., Koschate N., Totzek D. 2010 How price increases affect future purchases: The role of mental budgeting, income, and framing. J Psychology Marketing 27(1), 36–53. (doi:10.1002/mar.20318).

3. Bronner F., de Hoog R. 2016 Crisis resistance of tourist demand: the importance of quality of life. J Journal of travel research 55(2), 190–204. (doi:10.1177/0047287514541006).

4. Morewedge C.K., Holtzman L., Epley N. 2007 Unfixed Resources: Perceived Costs, Consumption, and the Accessible Account Effect. J Consum Res 34(4), 459–467. (doi:10.1086/518540).

5. van Wingerden M., Marx C., Kalenscher T. 2015 Budget Constraints Affect Male Rats’ Choices between Differently Priced Commodities. PLoS One 10(6), e0129581. (doi:10.1371/journal.pone.0129581).

6. Kagel J.H., Battalio R.C., Rachlin H., Green L. 1981 Demand Curves for Animal Consumers. Q J Econ 96(1), 1–15. (doi:Doi 10.2307/2936137).

7. Kagel J.H., Battalio R.C., Green L. 1995 Economic choice theory: An experimental analysis of animal behavior, Cambridge University Press.

8. Hu Y., van Wingerden M., Sellitto M., Schable S., Kalenscher T. 2021 Anterior Cingulate Cortex Lesions Abolish Budget Effects on Effort-Based Decision-Making in Rat Consumers. J Neurosci 41(20), 4448–4460. (doi:10.1523/JNEUROSCI.2541-20.2021).

9. Chen M.K., Lakshminarayanan V., Santos L.R. 2006 How basic are behavioral biases? Evidence from capuchin monkey trading behavior. J Polit Econ 114(3), 517–537. (doi:Doi 10.1086/503550).

10. Barron H.C., Mars R.B., Dupret D., Lerch J.P., Sampaio-Baptista C. 2021 Cross-species neuroscience: closing the explanatory gap. J Philosophical Transactions of the Royal Society B 376(1815), 20190633. (doi:10.1098/rstb.2019.0633).

11. Kalenscher T., van Wingerden M. 2011 Why we should use animals to study economic decision making - a perspective. Front Neurosci-Switz 5(82). (doi:10.3389/fnins.2011.00082).

12. Keysers C., Gazzola V. 2017 A Plea for Cross-species Social Neuroscience. Curr Top Behav Neurosci 30, 179–191. (doi:10.1007/7854_2016_439).

13. Garcia B., Cerrotti F., Palminteri S. 2021 The description-experience gap: a challenge for the neuroeconomics of decision-making under uncertainty. Philos Trans R Soc Lond B Biol Sci 376(1819), 20190665. (doi:10.1098/rstb.2019.0665).

14. Addessi E., Beran M.J., Bourgeois-Gironde S., Brosnan S.F., Leca J.B. 2020 Are the roots of human economic systems shared with non-human primates? Neurosci Biobehav Rev 109, 1–15. (doi:10.1016/j.neubiorev.2019.12.026).

15. McClure S.M., Ericson K.M., Laibson D.I., Loewenstein G., Cohen J.D. 2007 Time discounting for primary rewards. Journal of Neuroscience 27(21), 5796–5804. (doi:10.1523/Jneurosci.4246-06.2007).

16. Prevost C., Pessiglione M., Metereau E., Clery-Melin M.L., Dreher J.C. 2010 Separate valuation subsystems for delay and effort decision costs. J Neurosci 30(42), 14080–14090. (doi:10.1523/JNEUROSCI.2752-10.2010).

17. Klein-Flugge M.C., Kennerley S.W., Saraiva A.C., Penny W.D., Bestmann S. 2015 Behavioral Modeling of Human Choices Reveals Dissociable Effects of Physical Effort and Temporal Delay on Reward Devaluation. Plos Computational Biology 11(3). (doi:10.1371/journal.pcbi.1004116).

18. Croxson P.L., Walton M.E., O’Reilly J.X., Behrens T.E., Rushworth M.F. 2009 Effort-based cost-benefit valuation and the human brain. J Neurosci 29(14), 4531–4541. (doi:10.1523/JNEUROSCI.4515-08.2009).

19. Crockett M.J., Braams B.R., Clark L., Tobler P.N., Robbins T.W., Kalenscher T. 2013 Restricting temptations: neural mechanisms of precommitment. Neuron 79(2), 391–401. (doi:10.1016/j.neuron.2013.05.028).

20. Wilson M., Daly M. 2004 Do pretty women inspire men to discount the future? Proceedings of the Royal Society of London Series B: Biological Sciences 271(suppl_4), S177–S179. (doi:10.1098/rsbl.2003.0134).

21. Archambault C., Kalenscher T., de Laat J.J.J.o.B.D.M. 2020 Generosity and livelihoods: Dictator game evidence on the multidimensional nature of sharing among the Kenyan Maasai. 33(2), 196–207.

22. Ma Q., Hu Y. 2015 Beauty matters: social preferences in a three-person ultimatum game. PLoS One 10(5), e0125806. (doi:10.1371/journal.pone.0125806).

23. Otterbring T., Sela Y. 2020 Sexually arousing ads induce sex-specific financial decisions in hungry individuals. Personality and Individual Differences 152. (doi:10.1016/j.paid.2019.109576).

24. Li X.P., Zhang M. 2014 The Effects of Heightened Physiological Needs on Perception of Psychological Connectedness. J Consum Res 41(4), 1078–1088. (doi:10.1086/678051).

25. De Ridder D., Kroese F., Adriaanse M., Evers C. 2014 Always gamble on an empty stomach: Hunger is associated with advantageous decision making. PloS one 9(10), e111081. (doi:10.1371/journal.pone.0111081).

26. Ariely D., Loewenstein G. 2006 The heat of the moment: The effect of sexual arousal on sexual decision making. J Behav Decis Making 19(2), 87–98. (doi:10.1002/bdm.501).

27. Sescousse G., Barbalat G., Domenech P., Dreher J.C. 2013 Imbalance in the sensitivity to different types of rewards in pathological gambling. Brain 136, 2527–2538. (doi:10.1093/brain/awt126).

28. Jahedi S., Deck C., Ariely D. 2017 Arousal and economic decision making. Journal of Economic Behavior & Organization 134, 165–189. (doi:10.1016/j.jebo.2016.10.008).

29. Girard R., Obeso I., Thobois S., Park S.A., Vidal T., Favre E., Ulla M., Broussolle E., Krack P., Durif F., et al. 2019 Wait and you shall see: sexual delay discounting in hypersexual Parkinsons disease. Brain 142, 146–162. (doi:10.1093/brain/awy298).

30. Slutsky E.E., Ragusa O. 2012 On the Theory of the Budget of the Consumer. Giornale degli Economisti e Annali di Economia 71(2/3), 173–200.

31. Studer B., Koch C., Knecht S., Kalenscher T. 2019 Conquering the inner couch potato: precommitment is an effective strategy to enhance motivation for effortful actions. Philosophical Transactions of the Royal Society B 374(1766). (doi:10.1098/rstb.2018.0131).

32. Kurniawan I.T., Guitart-Masip M., Dayan P., Dolan R.J. 2013 Effort and valuation in the brain: the effects of anticipation and execution. J Neurosci 33(14), 6160–6169. (doi:10.1523/JNEUROSCI.4777-12.2013).

33. Klein-Flugge M.C., Kennerley S.W., Friston K., Bestmann S. 2016 Neural Signatures of Value Comparison in Human Cingulate Cortex during Decisions Requiring an Effort-Reward Trade-off. J Neurosci 36(39), 10002–10015. (doi:10.1523/JNEUROSCI.0292-16.2016).

34. Afriat S.N. 1972 Efficiency Estimation of Production Functions. International Economic Review 13(3), 568–598. (doi:10.2307/2525845).

35. Afriat S.N. 1973 On a System of Inequalities in Demand Analysis: An Extension of the Classical Method. International Economic Review 14(2), 460–472. (doi:10.2307/2525934).

36. Varian H.R. 1982 The Nonparametric Approach to Demand Analysis. Econometrica 50(4), 945–973. (doi:10.2307/1912771).

37. Glimcher P.W. 2011 Foundations of neuroeconomic analysis, OUP USA.

38. Nitsch F.J., Kalenscher T. 2020 Keeping a cool head at all times. What determines choice consistency?

39. JASP Team J. 2020 JASP (Version 0.14.1)[Computer software]. (

40. Ripley B., Venables B., Bates D., Hornik K., Gebhardt A., Firth D., Ripley M. 2013 Package ‘mass’. Cran R 538. (

41. Allen M., Poggiali D., Whitaker K., Marshall T., Kievit R. 2019 Raincloud plots: a multi-platform tool for robust data visualization [version 1; peer review: 2 approved]. 4(63). (doi:10.12688/wellcomeopenres.15191.1).

42. Loewenstein G. 2000 Emotions in economic theory and economic behavior. Am Econ Rev 90(2), 426–432. (doi:Doi 10.1257/aer.90.2.426).

43. Loewenstein G. 1996 Out of control: Visceral influences on behavior. Organ Behav Hum Dec 65(3), 272–292. (doi:10.1006/obhd.1996.0028).

44. Kahneman D., Thaler R.H. 2006 Anomalies - Utility maximization and experienced utility. J Econ Perspect 20(1), 221–234. (doi:Doi 10.1257/089533006776526076).

45. Loewenstein G., O’Donoghue T., Rabin M. 2003 Projection bias in predicting future utility. Q J Econ 118(4), 1209–1248. (doi:10.1162/003355303322552784).

46. Sweis B.M., Thomas M.J., Redish A.D. 2018 Mice learn to avoid regret. PLoS Biol 16(6), e2005853. (doi:10.1371/journal.pbio.2005853).

47. Sweis B.M., Abram S.V., Schmidt B.J., Seeland K.D., MacDonald A.W., 3rd, Thomas M.J., Redish A.D. 2018 Sensitivity to “sunk costs” in mice, rats, and humans. Science 361(6398), 178–181. (doi:10.1126/science.aar8644).

48. Constantinople C.M., Piet A.T., Brody C.D. 2019 An Analysis of Decision under Risk in Rats. Current Biology 29(12), 2066–2074. (doi:10.1016/j.cub.2019.05.013).

49. Oberliessen L., Hernandez-Lallement J., Schable S., van Wingerden M., Seinstra M., Kalenscher T. 2016 Inequity aversion in rats, Rattus norvegicus. Anim Behav 115, 157–166. (doi:10.1016/j.anbehav.2016.03.007).

50. Rudebeck P.H., Walton M.E., Smyth A.N., Bannerman D.M., Rushworth M.F. 2006 Separate neural pathways process different decision costs. Nat Neurosci 9(9), 1161–1168. (doi:10.1038/nn1756).

51. Salamone J.D., Correa M. 2018 Neurobiology and pharmacology of activational and effort-related aspects of motivation: rodent studies. Current Opinion in Behavioral Sciences 22, 114–120. (doi:10.1016/j.cobeha.2018.01.026).

52. Erev I., Roth A.E. 2014 Maximization, learning, and economic behavior. Proceedings of the National Academy of Sciences 111(Supplement 3), 10818–10825. (doi:10.1073/pnas.1402846111).

53. Monteiro T., Vasconcelos M., Kacelnik A. 2020 Choosing fast and simply: Construction of preferences by starlings through parallel option valuation. Plos Biology 18(8). (doi:10.1371/journal.pbio.3000841).

54. Lukinova E., Wang Y.Y., Lehrer S.F., Erlich J.C. 2019 Time preferences are reliable across time-horizons and verbal versus experiential tasks. Elife 8. (doi:10.7554/eLife.39656).

55. Hertwig R., Erev I. 2009 The description–experience gap in risky choice. Trends in Cognitive Sciences 13(12), 517–523. (doi:10.1016/j.tics.2009.09.004).

56. Heilbronner S.R., Hayden B.Y. 2016 The description-experience gap in risky choice in nonhuman primates. Psychon Bull Rev 23(2), 593–600. (doi:10.3758/s13423-015-0924-2).

57. Epley N., Mak D., Idson L.C. 2006 Bonus of rebate: the impact of income framing on spending and saving. J Behav Decis Making 19(3), 213–227. (doi:10.1002/bdm.519).

58. Shah A.K., Shafir E., Mullainathan S. 2015 Scarcity frames value. Psychol Sci 26(4), 402–412. (doi:10.1177/0956797614563958).

59. Ulkumen G., Thomas M., Morwitz V.G. 2008 Will I spend more in 12 months or a year? The effect of ease of estimation and confidence on budget estimates. J Consum Res 35(2), 245–256. (doi:10.1086/587627).

60. Stilley K.M., Inman J.J., Wakefield K.L. 2010 Planning to Make Unplanned Purchases? The Role of In-Store Slack in Budget Deviation. J Consum Res 37(2), 264–278. (doi:10.1086/651567).

61. Economides M., Guitart-Masip M., Kurth-Nelson Z., Dolan R.J. 2014 Anterior cingulate cortex instigates adaptive switches in choice by integrating immediate and delayed components of value in ventromedial prefrontal cortex. J Neurosci 34(9), 3340–3349. (doi:10.1523/JNEUROSCI.4313-13.2014).

62. Seinstra M.S., Sellitto M., Kalenscher T. 2018 Rate maximization and hyperbolic discounting in human experiential intertemporal decision making. Behav Ecol 29(1), 193–203. (doi:10.1093/beheco/arx145).

63. Constantino S.M., Daw N.D. 2015 Learning the opportunity cost of time in a patch-foraging task. Cogn Affect Behav Neurosci 15(4), 837–853. (doi:10.3758/s13415-015-0350-y).

64. Camille N., Griffiths C.A., Vo K., Fellows L.K., Kable J.W. 2011 Ventromedial frontal lobe damage disrupts value maximization in humans. J Neurosci 31(20), 7527–7532. (doi:10.1523/JNEUROSCI.6527-10.2011).

65. Chung H.K., Tymula A., Glimcher P. 2017 The Reduction of Ventrolateral Prefrontal Cortex Gray Matter Volume Correlates with Loss of Economic Rationality in Aging. Journal of Neuroscience 37(49), 12068–12077. (doi:10.1523/Jneurosci.1171-17.2017).

66. Harbaugh W.T., Krause K., Berry T.R. 2001 GARP for kids: On the development of rational choice behavior. Am Econ Rev 91(5), 1539–1545. (doi:Doi 10.1257/aer.91.5.1539).

67. Weinrabe A., Chung H.K., Tymula A., Tran J., Hickie I.B. 2020 Economic Rationality in Youth With Emerging Mood Disorders. J Neurosci Psychol E 13(3), 164–177. (doi:10.1037/npe0000129).

68. Kaufman B.E. 1999 Emotional arousal as a source of bounded rationality. Journal of Economic Behavior & Organization 38(2), 135–144. (doi:10.1016/S0167-2681(99)00002-5).

69. Pham M.T. 2007 Emotion and rationality: A critical review and interpretation of empirical evidence. Rev Gen Psychol 11(2), 155–178. (doi:10.1037/1089-2680.11.2.155).

70. Kurniawan I.T., Grueschow M., Ruff C.C. 2021 Anticipatory energization revealed by pupil and brain activity guides human effort-based decision making. J Neurosci. (doi:10.1523/JNEUROSCI.3027-20.2021).

