## Supplementary material for "Cross-species comparison of human and rodent primary reward consumption under budget constraints": Electronic Supplementary Material

**Supplementary Methods**

***Control Experiment 1***

To test the potential influence of reward satiation on milk consumptions, we increased the exchange ratio of effort-milk trade-off in the milk task of Control Experiment 1 (n = 16, mean age = 23, SD = 2) in which for each unit of purchase participants would obtain a doubled amount (i.e. 2ml) of milks as compared to the milk task of the main experiment (i.e. 1ml for each unit), while the picture task remained the same. All other procedures and measures were the same as the milk and picture task of human experiment reported in the main text.

***Control Experiment 2***

It is possible that the violations of normative rationality in the picture task are mostly driven by the mode of reward delivery at the end of each session since the effect of viewing a continuous presentation of pictures might be more satisfying or arousing than viewing a single unit of stimuli on a trial-by-trial basis. To control for this possibility, we included Control Experiment 2 (n = 25, mean age = 23, SD = 2) where picture rewards were delivered and consumed in a trial-by trial manner (Figure S1-E). There were two modifications in the trial structure as compared to the main experiment and the first control experiment: (i) picture rewards were delivered and consumed in a trial-by-trial manner, i.e., participants viewed a photo of the selected category of picture rewards immediately after their effort exertion phase; (ii) participants received no updates on their selected choice bundle within the session. The block design and price ratio (i.e. 4:4 PP for baselines, 8:2 PP for budget conditions) remained the same, except that participants had a doubled amount of budget to spend in each experimental condition. The baseline and uncompensated budget were 160 PPs and individual compensated budget was extended according to the Slutsky equation.

***Rat Experiment***

*Subjects and Rewards*

Batch 1 (n = 7) received saline injection in the nucleus accumbens and batch 2 (n = 10) received saline injection in the medial orbitofrontal cortex. Histological verification after the experiments confirmed that no tissue damage occurred in the respective brain regions. All animals weighed between 280 - 350g at the beginning of the training and had not been used in any previous testing. Rats were housed in a group of three and under a reversed 12:12h light-dark cycle, at a temperature of 22 ± 2 ºC and humidity-controlled (60%) colony room. Throughout the experiment, all animals were food deprived with water available ad libitum in the home cages, and they were kept at 85% of free-feeding body weight.

***Simulation***

Though we’ve found consistent behaviors in rats across batches and experiments, one could argue against the validity of cross-species comparison given the difference in the sample size of human and animal experiment. To examine whether the comparability and discrepancy in the cross-species behaviors were merely driven by a sampling bias [1, 2], we generated n = 1000 choice sets (i.e. choice bundles) following a uniform distribution. Each choice set contained five choice bundles for the five experimental conditions of human budget task. Each choice bundle is referred to a combination of the quantity of two rewards ($q_{P}, q_{NP}),$calculated from equation (1), given the fixed budgets (*m*) and price regimes of the three baselines (${price}_{P}$: ${price}_{NP}$ = 4:4) and the two budget conditions (${price}_{P}$: ${price}_{NP}$ = 8:2):

${price}_{P}{\times q}_{P} +{price}_{NP}{\times q}_{NP} =m$	(1)

from which we defined the preferred and non-preferred reward based on the preceding baseline choice and calculated the demand elasticity of each reward commodity in the uncompensated and compensated budget condition.

***Analysis***

Elasticity data were analyzed in JASP using repeated measures ANOVA with Bonferroni correction for multiple post-hoc comparisons. Wilcoxon signed rank test was performed in R for comparing self-reported score of state before and after each task. The linear modeling on the elasticity of non-preferred rewards against scaled baseline choice of preferred rewards and the budget conditions (i.e. uncompensated and compensated) was performed with a general Gaussian linear regression.

**Supplementary Results**

**Control Experiment 1: double the amount of milk per unit did not affect participant’s elasticity pattern of milk rewards**

As expected (Figure S1-B), participants also reported a decrease in hunger after the milk task (p = 0.008, Wilcoxon signed-rank test), and an increased sexual desire after the picture task (p = 0.006, Wilcoxon signed-rank test). We conducted a similar three-way ANOVA with budget condition (uncompensated, compensated), task (milk, picture) and preference (preferred commodity, non-preferred commodity) on reward elasticities (Figure S1-A). We found a main effect of budget (F_1,15_ = 10.753, p = 0.005, partial η^2^ = 0.418) and a marginally significant main effect of task (F_1,15_ = 4.347, p = 0.055, partial η^2^ = 0.225). We also found a significant *budget x preference* interaction (F_1,15_ = 8.827, p = 0.01, partial η^2^ = 0.37) and a marginally significant interaction of *preference x task* on demand elasticity (F_1,15_ = 3.998, p = 0.064, partial η^2^ = 0.21). The three-way interaction of *budget x preference x task* was not significant (F_1,15_ = 0.197, p = 0.664, partial η^2^ = 0.013). Therefore, we further performed two separate 2 (budget: compensated vs uncompensated) x 2 (preference: preferred vs non-preferred) ANOVA on reward elasticities for the milk and picture task of Control Experiment 1.

In the milk task, we found a main effect of budget (F_1,15_ = 6.318, p = 0.024, partial η^2^ = 0.296), a marginally significant main effect of preference (F_1,15_ = 4.3, p = 0.056, partial η^2^ = 0.223) and an interaction of *budget x preference* on reward elasticities (F_1,15_ = 6.341, p = 0.024, partial η^2^ = 0.297). Consistent with the pattern observed in the milk task (1ml version) of human and rat experiment, we found a null difference between the two reward elasticities in the uncompensated budget condition (t = 0.338, p = 0.74, Cohen’s d = 0.084) which was enlarged in the compensated budget condition (t = 3.098, p < 0.001, Cohen’s d = 0.775). Having an extended budget also reduced the negativity of demand elasticity for preferred milk rewards (t = -4.616, p < 0.001, Cohen’s d = -1.154), but did not influence the elasticity of non-preferred rewards (t = -0.568, p = 0.579, Cohen’s d = -0.142). For the picture task, the interaction of *budget x preference* revealed a tendency towards significance (F_1,15_ = 3.817, p = 0.07, partial η^2^ = 0.203). We found no main effect of budget (F_1,15_ = 1.771, p = 0.203, partial η^2^ = 0.106), nor preference (F_1,15_ = 0.316, p = 0.582, partial η^2^ = 0.021) on reward elasticities. We interpret these findings as a replication of the milk task in the main experiment that increasing the value of single unit of milk rewards, as well as the total amount of milk intake did not alter individual’s consumption pattern. Worth noting is, when controlling the sample size in this control experiment (n = 16) that was more comparable with the rat experiment (n = 17), humans revealed a consistent larger variance in the population than rats in utilizing effort budget to trade for rewards.

**Control Experiment 2: timing of reward consumption did not affect participants’ elasticity pattern of picture rewards**

We performed a two-way ANOVA with budget (uncompensated, compensated) and preference (preferred commodity, non-preferred commodity) on demand elasticity of picture rewards. We found a main effect of preference (F_1,24_ = 8.137, p = 0.009, partial η^2^ = 0.253) and an interaction of *budget x preference* (F_1,24_ = 9.802, p = 0.005, partial η^2^ = 0.29) on reward elasticities (Figure S1-C). Importantly, we replicated our finding in the picture task of the main experiment that the elasticity of non-preferred rewards in the uncompensated budget condition was much more positive than the elasticity of preferred rewards (t = -4.733, p < 0.001, Cohen’s d = -0.947). Such comparison was not significant in the compensated budget condition (t = -0.578, p = 0.569, Cohen’s d = -0.116). We also found a reduced negative elasticity of preferred reward in the compensated budget condition as compared to the uncompensated condition (t = -11.077, p < 0.001, Cohen’s d = -2.215). The elasticity of non-preferred rewards did not differ between the two budget conditions (t = 1.29, p = 0.209, Cohen’s d = 0.258). Also, we found a tendency on participants of Control Experiment 2 showing an adaptative reward selection in the compensated budget condition (approaching significance, t = -1.954, p = 0.063), but not in the uncompensated condition (t = -0.757, p = 0.459, Figure S1-D).

Put together, we found that increasing the value of single units of milk, as well as the total amount of milk intake did not alter the individuals’ consumption pattern of milk rewards. Also, the timing and manner of reward consumption did not affect participants’ choice pattern of picture rewards. Thus, we concluded that our findings of the distinct strategy of budget and choice allocation regarding the two types of primary rewards to be a robust and replicable behavioral phenomenon on human participants.

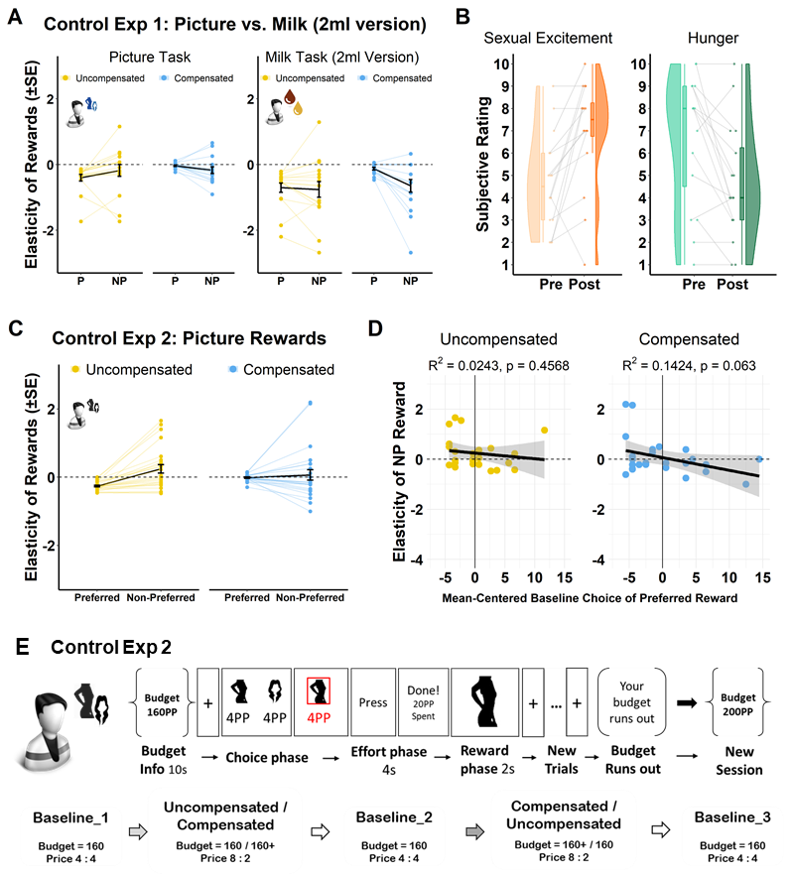
***Figure S1 Two control experiments on the effect of reward satiation and timing of reward delivery on consumption of milk and picture rewards.*** *(A-B) Control Experiment 1: demand elasticity of rewards and self-reported scores of sexual desire and hunger before (“Pre”) and after (“Post”) the task. Each unit of purchase in the milk task would yield the doubled amount (i.e. 2ml) of milk compared to the milk task of main experiment (i.e. 1ml for each unit), while the picture task remained the same. (C-D) Control Experiment 2: elasticity of preferred and non-preferred picture rewards and correlations between the scaled baseline choice of preferred rewards and elasticity of non-preferred rewards in the two budget conditions. (E) Schematic representation of Control Experiment 2 in which picture rewards were delivered and consumed at the end of each trial, that participants viewed a photo of the selected category of picture rewards immediately after their effort exertion phase. P: preferred rewards; NP: non-preferred rewards.*

**Further Supplemental Results**

**Comparable baseline-preference-dependent reward substitution in rats and humans in the milk task, not in the picture task**

In the main text, we’ve shown a comparable elasticity pattern in rats and humans when spending a limited budget to obtain milk rewards. However, it remains unclear whether humans and rats underwent the same decision process that led to the approximation in their consumption pattern in the budget conditions. Our previous study [3] has shown that the extent of rats’ substituting cheap vanilla milk with expensive preferred chocolate in the uncompensated budget condition, indicated by the elasticity of vanilla, was dependent on the strength of baseline preference for chocolate. This revealed a strategy adaptation in the sham rats when experiencing the shrinking purchasing power of the uncompensated budget. If humans adopted a similar strategy to adapt their reward selection after the price change, the elasticity of non-preferred reward should be linked with individual baseline preference in the milk task, too, but not in the picture task where they held a rigid preference for priced picture rewards. Furthermore, we expected such null adaptation in the picture task to be prominent in the uncompensated budget condition, but not in the compensated condition where individual budget got extended to restore the purchasing power of the priced commodity. To test this hypothesis, we regressed the elasticity of non-preferred rewards against the scaled baseline choice of preferred rewards, the budget condition and the interaction between the two, for both rats and humans.

In line with our previous study, we found a main effect of baseline choice of preferred reward (t = -3.843, p < 0.001) on the elasticity of non-preferred reward in rats (Figure S2). Neither the main effect of budget condition (t = -0.279, p = 0.782) nor interaction between the two (t = -0.181, p = 0.857) was significant. Specifically, the association between the elasticity of non-preferred rewards and the baseline preference was significant for the uncompensated budget condition (t = -3.832, p = 0.002), and for the compensated budget condition (t = -6.741, p < 0.001). For the human milk task (Figure S2), we found a main effect of the baseline preference (t = -3.766, p < 0.001) as well as a main effect of budget condition (t = 1.986, p = 0.049) on the elasticity of non-preferred reward, but no significant interaction of the two (t = 0.19, p = 0.849). Similarly, the elasticity of non-preferred rewards was negatively correlated with the scaled baseline choice of preferred rewards in the two budget conditions of the milk task (uncompensated: t = -3.605, p < 0.001; compensated: t = -3.853, p < 0.001). For the human picture task (Figure S2), as predicted, we found a strong correlation between the elasticity of non-preferred rewards and baseline preference in the compensated budget condition of the picture task (t = -3.593, p < 0.001), but not in the uncompensated budget condition (t = -0.92, p = 0.362). The main effect of baseline choice of preferred rewards (t = -3.577, p < 0.001) and budget condition (t = 2.317, p = 0.022) were significant in the picture task. The interaction between the two was near-significant (t = 1.842, p = 0.068).

To conclude, these replicate the baseline-preference-dependent reward substitution in the sham rats and reveal a similar adaptive process in humans in the milk task. In another words, when humans and rats experienced a price shock of preferred milk rewards, the pressure of deploying an expensive-to-cheap shift was not equal among individuals, but closely linked to their baseline consumption level. For those who held a higher baseline preference for the preferred milks, they also shifted more heavily from preferred to non-preferred milk reward after the price change. Such behavioral adaptation was also present in humans in the picture task when the individual budget was extended to allow the reselection of baseline choice bundle, but not in the uncompensated budget condition where participants showed a larger resistance to the pressure of price increase when the spendable resource was in scare, thus, led to the positive elasticity of picture rewards.

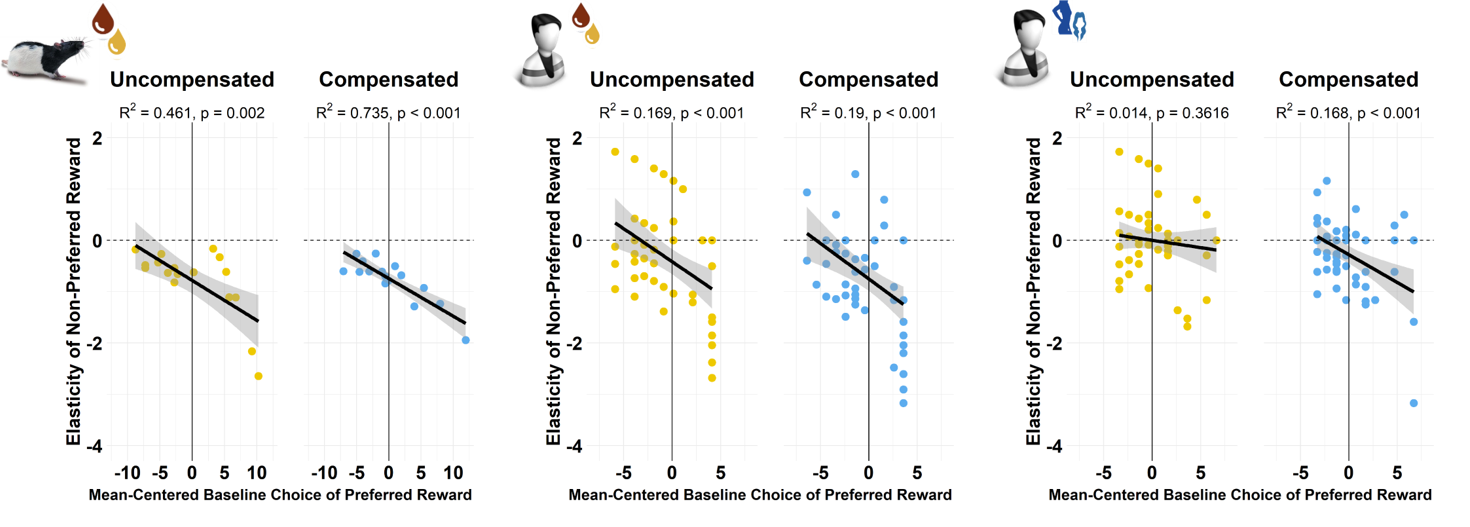
**Figure S2 Baseline-preference-dependent reward substitution in rats (left), the milk task (middle) but not in the picture task (right).** *We regressed the elasticity of non-preferred rewards against the scaled baseline choices of preferred rewards, the budget condition and the interaction between the two, for both rats and humans.*

**Distribution of individual reward substitutions of rats and humans in the Elasticity Matrix**

As reasoned above, to examine whether there existed a sampling bias in the rat experiment, we plotted the elasticities of the two reward commodities on a “Elasticity Matrix” for the rats, humans and simulated data (Figure S3, see details in the *Supplementary Method*). The position of individual data on the matrix indicates the level and direction of reward substitution after the price change. In the uncompensated budget condition, a majority of the simulated sample (854/1000) showed negative elasticities for both rewards, mainly distributed around the diagonal line in the third quadrant of the matrix, suggesting a relatively equal increase and decrease of demand for cheaper non-preferred and expensive preferred rewards. In the compensated condition, more simulated individuals (907/1000) showed negative elasticities for both rewards but drifted towards a reduced negativity of demand for preferred reward (i.e. a less reduction from the baseline demand) and an increased negative elasticity (i.e. a larger increase from the baseline demand) of non-preferred reward. Importantly, an approximate proportion of population in rats (17/17) exhibiting the price-sensitive budget-dependent consumption behavior, as well as in humans in the milk task (uncompensated: 47/60, compensated: 53/60). However, the distribution of individual elasticities in the human picture task shifted away from the third quadrant and more often located in the second (25/60) and first quadrant (16/60) of the matrix. These choice sets were considered as a suboptimal decision strategy in response to the price change and budget constraints. These results speak to the little influence of sample size difference on running a cross-species comparison in our experiments. Rather, on the individual level, rat consumers exhibited a much less between-subject variability as compared to humans in the milk and picture task. It suggests that rigorous animal models can serve as a benchmark to evaluate humans’ economic decision making.

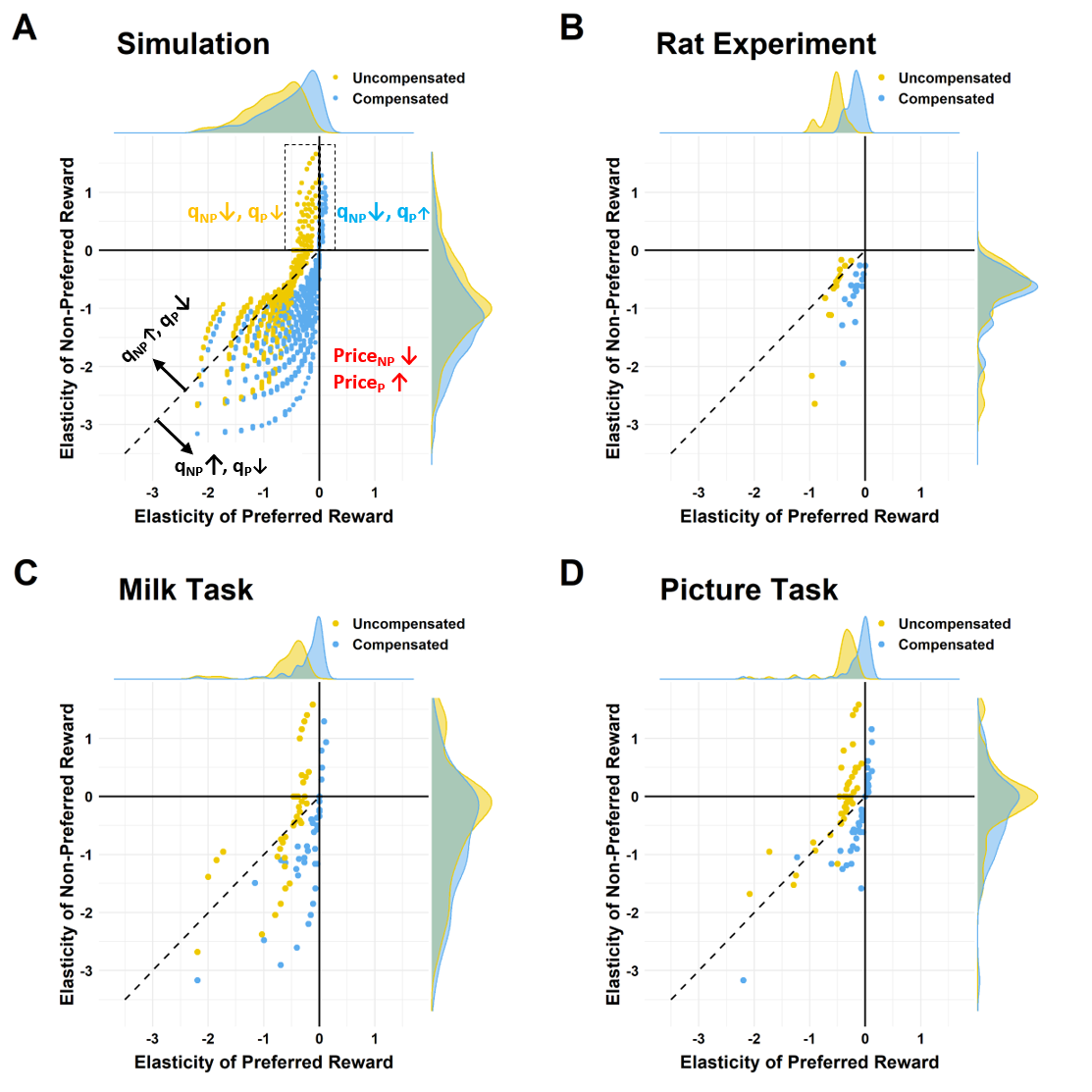
**Figure S3 *Elasticity Matrix:*** *demand elasticities of preferred and non-preferred rewards in the uncompensated and compensated budget condition of the simulated choice sets (A), Rat Experiment (B), milk task (C) and picture task (D) of Human Experiment. The position of individual data on the matrix indicates the level and direction of reward substitution after the price change. Dots represent each individual data; on the upper and right side of the matrix is the density plot of the distribution; the dash line indicates the diagonal line (y = x). Price_P_: price of preferred reward; Price_NP_: price of non-preferred reward; q_P_: quantity of preferred reward; q_NP_: quantity of non-preferred reward. Upward arrow indicates an increase in the price or quantity of reward choice, downward arrow indicates a decrease in the price or quantity of reward choice.*

**Additional ANOVA result of *preference x task* on reward elasticity in the human experiment**

As reported in the main text, besides the budget regulation on reward elasticities in the picture and milk task (Figure 2 D-E), we also found a significant interaction of *preference* *x task* on demand elasticity of rewards (F_1,59_ = 14.668, p < 0.001, partial η^2^ = 0.199). Breaking it down (Figure S4-A), the elasticity of preferred rewards, averaged for the two budget conditions, was less negative than the mean elasticity of non-preferred rewards in the milk task (t = 2.426, p = 0.018, Cohen’s d = 0.313), but was more negative than the non-preferred rewards in the picture task (t = -2.419, p = 0.019, Cohen’s d = -0.312). Also, the mean elasticity of preferred (t = -2.776, p = 0.027, Cohen’s d = -0.294) and non-preferred (t = -3.637, p < 0.001, Cohen’s d = -0.47) rewards in the milk task were much more negative than these two in the picture task. Importantly, we found a strong link between individual elasticity pattern and decision rationality in that the difference between the mean elasticity of preferred and non-preferred rewards was significantly correlated to the level of choice consistency indexed by the GARP violation count in the picture (z = -3.489, p < 0.001) and milk (z = -2.887, p = 0.004) task (Figure S4-A). Briefly speaking, participants who exhibited a more positive elasticity for non-preferred rewards, as compared to their elasticity of preferred rewards, showed a higher level of GARP violations. Such an elasticity-rationality correlation verified that our rationality test, despite its limited power given only five choice bundles, was not a random result but can truly reveal individual choice patterns.

**Intra-individual stability in the reward elasticity of picture and milk task**

Lastly, to explore whether there exists any intra-individual stability in the consumption strategy across the two tasks, we regressed the elasticity of non-preferred rewards in the uncompensated budget condition of picture task against that in the milk task and found a significant correlation between the two (t = 2.547, p = 0.014, Figure S4-B). It suggested that participants with a positive elasticity of non-preferred rewards in the picture task tended to be resistant in the choice adaptation to the price changes in the milk task when the budget constraint was high (i.e. uncompensated budget condition). This finding adds to the existing evidence revealing intra-individual stability in cost-benefit decision makings [4-6].

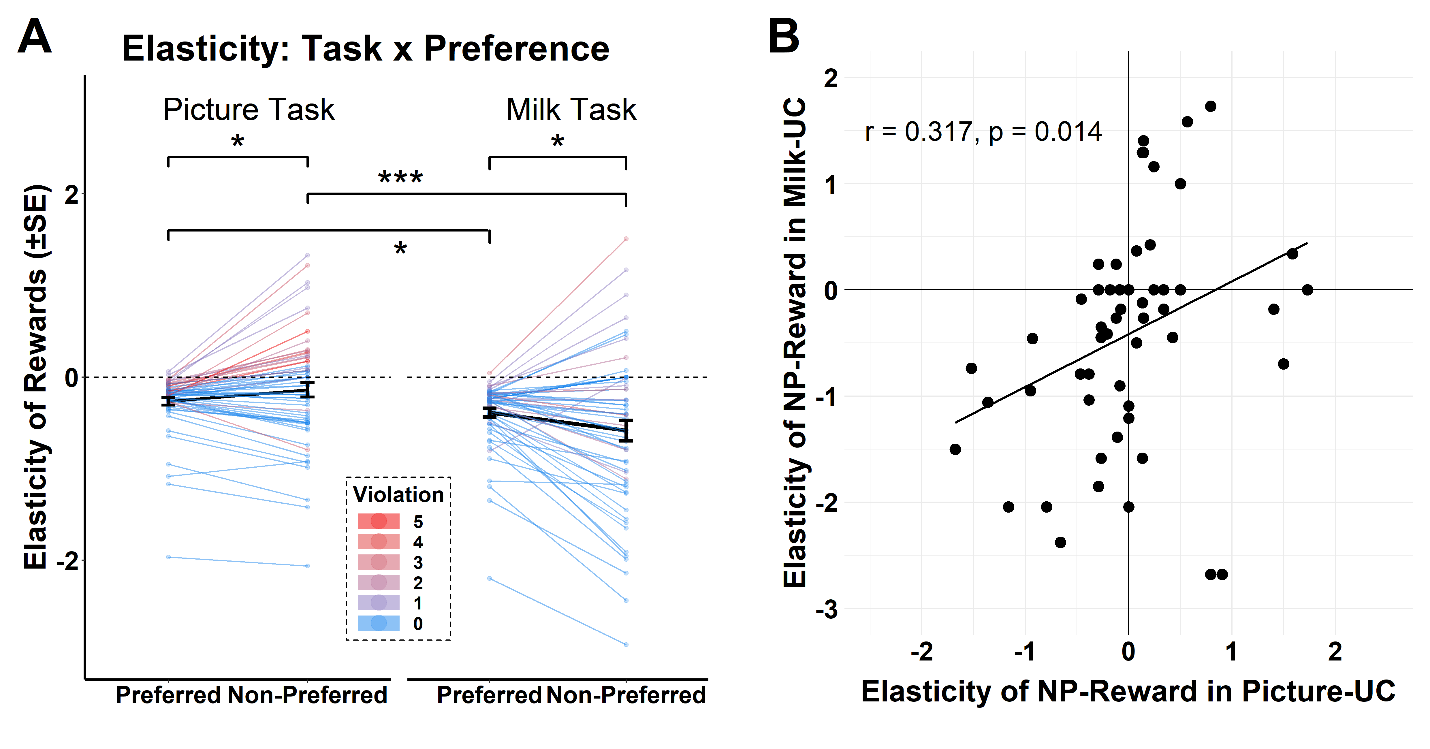
**Figure S4** *(A) Mean elasticity of preferred and non-preferred rewards in the two tasks, plotted against the count of GARP violation in each task. (B) Intra-individual stability in the reward elasticity of picture and milk task: correlation of non-preferred reward elasticity in the uncompensated budget condition in the picture and milk task.* *NP-Reward: non-preferred reward;* *UC: uncompensated budget condition. ***: p < 0.001, *: p < 0.05.*

**Reference**

1.	Stuber E.F., Araya-Ajoy Y.G., Mathot K.J., Mutzel A., Nicolaus M., Wijmenga J.J., Mueller J.C., Dingemanse N.J. 2013 Slow explorers take less risk: a problem of sampling bias in ecological studies. *Behav Ecol* **24**(5), 1092-1098. (doi:10.1093/beheco/art035).

2.	Michelangeli M., Wong B.B.M., Chapple D.G. 2015 It’s a trap: sampling bias due to animal personality is not always inevitable. *Behav Ecol* **27**(1), 62-67. (doi:10.1093/beheco/arv123).

3.	Hu Y., van Wingerden M., Sellitto M., Schable S., Kalenscher T. 2021 Anterior Cingulate Cortex Lesions Abolish Budget Effects on Effort-Based Decision-Making in Rat Consumers. *J Neurosci* **41**(20), 4448-4460. (doi:10.1523/JNEUROSCI.2541-20.2021).

4.	Reuben E., Sapienza P., Zingales L. 2010 Time discounting for primary and monetary rewards. *Econ Lett* **106**(2), 125-127. (doi:10.1016/j.econlet.2009.10.020).

5.	Studer B., Koch C., Knecht S., Kalenscher T. 2019 Conquering the inner couch potato: precommitment is an effective strategy to enhance motivation for effortful actions. *Philosophical Transactions of the Royal Society B* **374**(1766). (doi:10.1098/rstb.2018.0131).

6.	Lukinova E., Wang Y.Y., Lehrer S.F., Erlich J.C. 2019 Time preferences are reliable across time-horizons and verbal versus experiential tasks. *Elife* **8**. (doi:10.7554/eLife.39656).
